# SARS-CoV-2 S1 Subunit Booster Vaccination Elicits Robust Humoral Immune Responses in Aged Mice

**DOI:** 10.1101/2022.10.25.513090

**Authors:** Eun Kim, Muhammad S. Khan, Alessandro Ferrari, Shaohua Huang, Josè C. Sammartino, Elena Percivalle, Thomas W. Kenniston, Irene Cassaniti, Fausto Baldanti, Andrea Gambotto

**Author notes:** **Corresponding Author** Andrea Gambotto, MD, Department of Surgery, University of Pittsburgh School of Medicine, W1148 Biomedical Science Tower, 200 Lothrop St. Pittsburgh, PA 15261, USA.

## Abstract

Currently approved COVID-19 vaccines prevent symptomatic infection, hospitalization, and death of the disease. However, the emergence of severe acute respiratory syndrome coronavirus 2 (SARS-CoV-2) variants raises concerns of reduced vaccine effectiveness and increased risk of infection. Repeated homologous booster in elderly individuals and immunocompromised patients is considered to solve severe form of disease caused by new SARS-CoV-2 variants but cannot protect completely against breakthrough infection. In our previous study we assessed the immunogenicity of an adenovirus-based vaccine expressing SARS-CoV-2-S1 (Ad5.S1) in mice, resulting in that a single immunization with Ad5.S1, via subcutaneously injection or intranasal delivery, induced robust humoral and cellular immune responses [1]. As a follow up study, here we showed that vaccinated mice had high titers of anti-S1 antibodies at one year after vaccination compared to PBS immunized mice. Furthermore, one booster dose of non-adjuvanted recombinant S1Beta (rS1Beta) subunit vaccine was effective in stimulating strong long-lived S1-specific immune responses and inducing significantly high neutralizing antibodies against the Wuhan, Beta, and Delta strain with 3.6- to 19.5-fold change increases. Importantly, the booster dose elicits cross-reactive antibody responses resulting in ACE2 binding inhibition against spike of SARS-CoV-2 variants (Wuhan, Alpha, Beta, Gamma, Delta, Zeta, Kappa, New York, India) as early as two-week post-boost injection, persisting over 28 weeks after a booster vaccination. Interestingly, levels of neutralizing antibodies were correlated with not only level of S1-binding IgG but also level of ACE2 inhibition in the before- and after-booster serum samples. Our findings show that S1 recombinant protein subunit vaccine candidate as a booster has potential to offer cross-neutralization against broad variants, and has important implications for vaccine control of new emerging breakthrough SARS-CoV-2 variants in elderly individuals primed with adenovirus-based vaccine like AZD1222 and Ad26.COV2.S.

## Introduction

Severe acute respiratory syndrome coronavirus-2 (SARS-CoV-2) was identified as the causative agent of COVID-19 in December 2019 and resulted in a pandemic of coronavirus disease 2019 (COVID-19). The COVID-19 pandemic has 622 million confirmed cases, 6.5 million reported deaths, and 12.7 billion vaccine doses administered worldwide (until October 12, 2022) [2]. Six vaccines for SARS-CoV-2 targeting the spike (S) protein (BNT162b2; AZD1222; Ad26.COV2.S; mRNA-1273; NVX-CoV2373; Ad5-nCoV) have been approved by World Health Organization (WHO) and have greatly reduced the rate of severe disease and death [3]. However, SARS-CoV-2 evolution gave rise to multiple variants including SARS-CoV-2 variants of concern (VOCs), such as Alpha (B.1.1.7), Beta (B.1.351), Gamma (P.1), Delta (B.1.617.2) and lastly Omicron (B.1.1.529), characterized by potential increased transmissibility, neutralizing antibody escape, and reduced effectiveness of vaccinations or antibody treatment [4].

It is clear that age alone is the most significant risk factor for death due to COVID-19 [5–7]. Recent reports suggested that patients over 65 are responsible for 80% of COVID-19 hospitalizations and suffer from a 20-fold higher COVID-19 fatality rate compared to those under 65 years old [8–10], with individuals aged 80 or more representing the group at greatest risk of severe COVID-19 [11]. Furthermore, elderly individuals induced poor neutralization, which could be explained by lower serum IgG level and lower somatic hypermutation in B cell selection, along with lower IL-2-producing CD4^+^Tcells help compared to younger individuals, and all of which are overcame by booster [12]. These data are in line with previous findings that immune responses in aged mice vaccinated with ChAdOx1 nCov-19 was lower than those in younger mice, which was overcome by booster dosing [13].

The entry of coronaviruses into host cells is mediated by interaction between the receptor binding domain (RBD) of the viral S protein and the host receptor, angiotensin-converting enzyme 2 (ACE2) through the upper and lower respiratory tracts [14, 15]. Neutralizing antibodies against SARS-CoV-2 are effective at blocking this interaction to prevent infection [16, 17]. Competitive immunoassay for quantifying inhibition of the spike-ACE2 interaction have been shown to be a high level of concordance with neutralizing test [18, 19]. VOCs have mutations or deletions in the spike protein, with some mutations occurring in the RBD, resulting in the highest resistance to vaccine-induced and infection-acquired immunity. In response to the rapid evolution of SARS-CoV-2, and the global circulation of VOC, the booster injection has been considered to protect from breakthrough infections of new emerging variants. Evaluation of booster immunization have been investigated in mice, non-human primates, and human [13, 20–23]. The findings suggested that the level of neutralizing antibodies was correlated with vaccine efficacy for both mRNA and adenovirus vectored vaccine, and likely potential efficacy after boosting [24–27]. Of note, ChAdOx1-mRNA vaccination was safe and enhanced immunogenicity compared to ChAdOx1-ChAdOx1 vaccination, highlighting that heterologous prime-boost regimens may offer immunological advantages to elicit the strong and long-lasting protection acquired with currently available adenovirus-based vaccines [28–30]. Overall, a heterologous booster administration has been considered as a solution to protect elderly people from a breakthrough infection of new emerging variants.

In our previous study we assessed the immunogenicity of adenoviral based vaccine expressing SARS-CoV-2-S1 (Ad5.S1) in mice [1]. Here we conducted the follow up study to assess long term persistence of immunogenicity and the booster effect of subunit vaccine in aged mice. For the subunit vaccine, recombinant protein S1 of SARS-CoV-2 Beta (B.1.351) (rS1Beta) was selected, because it showed the greatest breakthrough infections against the Wuhan-based vaccines [31, 32], before COVID-19 waves by Omicron variants, which was shown to cause even higher levels of vaccine escape lately. From the present study it was evaluated that vaccinated mice with Ad5.S1 had high titers of anti-S1 antibodies after one-year immunization compared to PBS immunized mice and a booster with rS1Beta subunit vaccine was effective in stimulating strong long-lived S1-specific immune responses and in inducing significantly high cross-neutralizing antibodies against SARS-CoV-2 variants.

## Results

### Construction and expression of recombinant proteins

To produce recombinant proteins of SARS-CoV-2-S1, pAd/S1Beta was generated by subcloning the codon-optimized SARS-CoV-2-S1Beta gene having C-tag into the shuttle vector, pAd (GenBank U62024) at SalI & NotI sites **(Fig. 1A)**. To determine whether rS1Beta proteins were expressed from the plasmid, Expi293 cells were transfected with pAd/S1Beta or pAd as a control. At 5 days after transfection, the supernatants of Expi293 cells were characterized by a sandwich ELISA using monoclonal antibodies pair against SARS-CoV-2 Wuhan (WU) **(Fig. 1B)** and Western blot analysis **(Fig. 1C)**. As shown in Figure 1B, the titer of recombinant rS1Beta proteins expressed in Expi293 cells was about 7.3 mg/L based on a standard of rS1WU and about 40.0 mg/L based on a standard of rS1Beta, while rS1Beta protein did not detected in the Expi293 cells transfected with control pAd. The rS1Beta protein was separated by a 10% SDS-PAGE and recognized by a polyclonal anti-spike of SARS-CoV-2 antibody at the expected glycosylated monomeric molecular weights of about 110 kDa under the denaturing reduced conditions, while no expression was detected in the mock-transfected cells **(Fig. 1C).** The purified rS1Beta protein using C-tagXL affinity matrix was determined by silver staining **(Fig. 1D).**

**Figure 1.**
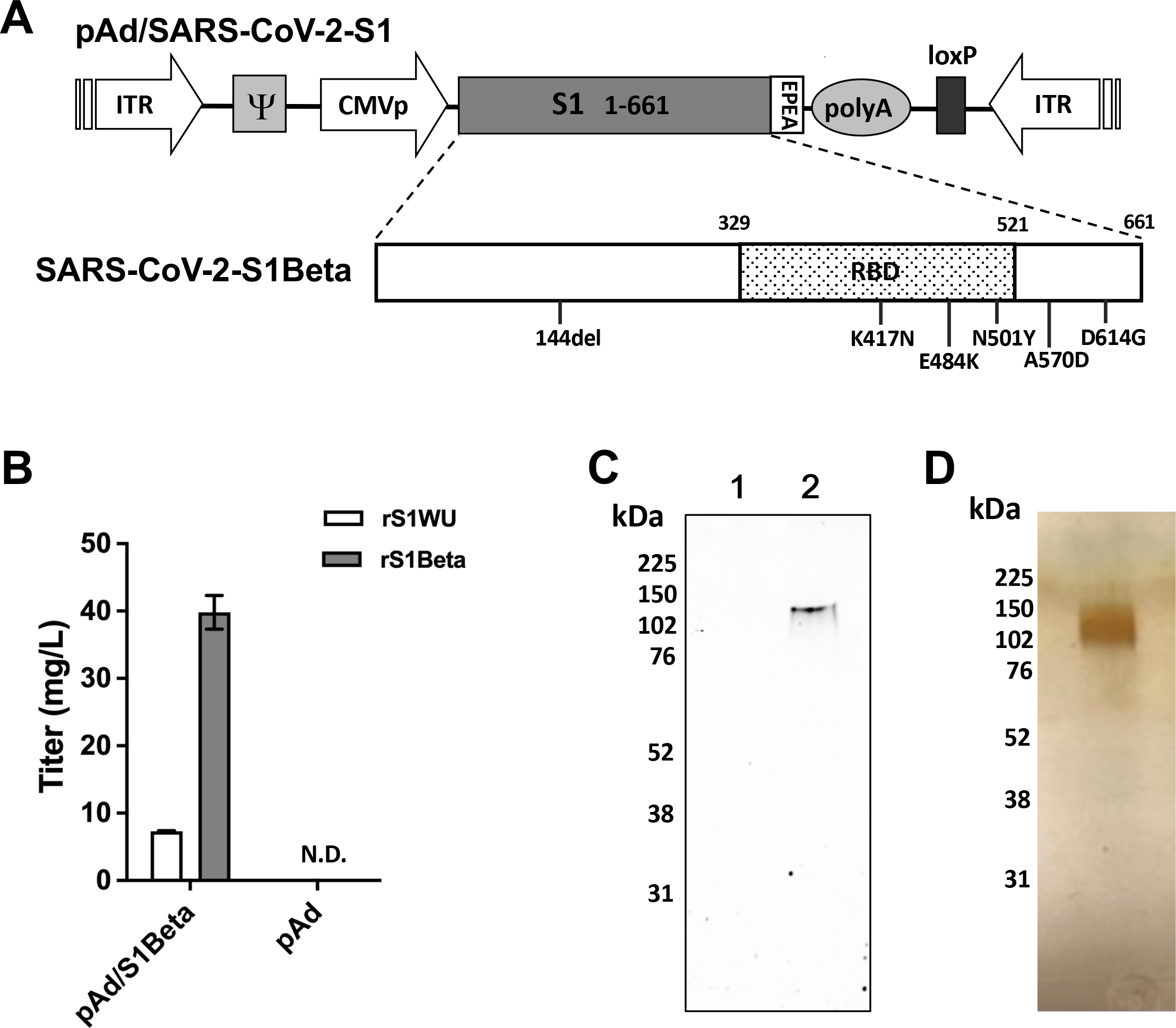
Construction of recombinant SARS-CoV-2-S1Beta protein expressing plasmid. **A.** A shuttle vector carrying the codon-optimized SARS-CoV-2-S1 gene of beta (1.351.1) variants encoding N-terminal 1-661 with c-tag (EPEA) was designated as shown in the diagram. Amino acid changes in the SARS-CoV-2-S1 region of in this study is shown. ITR: inverted terminal repeat; RBD: receptor binding domain. **B.** Titer of recombinant SARS-CoV-2-S1 proteins by sandwich ELISA with the supernatant of Expi293 cells transfected pAd/SARS-CoV-2-S1Beta (pAd/S1Beta) based on the standard of rS1Wuhan (WU) (white box) or rS1Beta (grey box) **C.** Detection of the SARS-CoV-2-S1 proteins by western blot with the supernatant of Expi293 cells transfected with pAd/S1Beta using anti spike protein of SARS-CoV-2 rabbit polyclonal antibody (lane 2). As a negative control, mock-transfected cells were treated the same (lane 1). The supernatants were resolved on SDS-10% polyacrylamide gel after being boiled in 2% SDS sample buffer with β-ME. **D.** Silver-stained reducing SDS-PAGE gel of purified Expi293 cell-derived rS1Beta (300ng).

### Rapid Recall of S1-Specific Binding Antibodies after a Booster

In our previous study we evaluated the immunogenicity of adenoviral vaccine until week 24 [1]. To assess long-term persistence of immunogenicity, we first determined antigen-specific IgG antibody endpoint titers in the sera of vaccinated mice (Ad5.S1 immunized groups either via I.N. delivery or S.C. injection) and control mice (PBS or Adψ5 immunized groups) at week 52, one year after vaccination. As shown in Figure 2A, significantly high titers of anti-S1 IgG antibodies were present in Ad5.S1 vaccinated mouse groups (G4, *p* = 0.0016 and G5, *p* = 0.0365) even after one year of vaccination as compared to AdΨ5-vaccinated mouse groups (G2 and G3) or PBS group (G1). To assess the booster effect of subunit vaccine, we collected serum samples from all mice before booster immunization (W52) and immunized animals with 15 μg of rS1Beta intramuscularly, collected sera subsequent weeks, and examined the end point titer of IgG against the S1 subunit of the spike protein (anti-S1) binding antibodies by ELISA **(Fig. 2A)**. We found more binding antibodies were detected significantly in Ad5.S1 vaccinated mouse groups (G4 and G5) compared to AdΨ5-vaccinated mouse groups (G2 and G3) or PBS group (G1) until week 80 (*p* < 0.05) after a booster vaccination. The change of geometric mean titers (GMT) of IgG end point titer in G4 and G5 compared to those at week 52 were same as 32-fold at week 54, and 55.7-fold and 18.4-fold at week 56, respectively **(Supplementary Fig.1A)**. Interestingly, the peak of IgG end point titer showed at week 56 (at week 4 after a booster) in G4, while it showed at week 54 (at week 2 after a booster) in G5. These recalls were faster after a booster vaccination with rS1Beta subunit vaccine when compared with IgG end point titer after prime (week 6 post-prime vs. week 2 or 4 post-boost) [1]. Furthermore, the elicited IgG antibody responses after a booster lasted longer, through week 80 (maximum length of the study to date), than after a prime, resulting from the comparison with IgG end point titers at week 28 post-prime (W28) or post-boost (W80) **(Supplementary Fig.1B)**. The mouse group primed subcutaneously (G4) had higher antibody titers as compared to mouse group primed intranasally (G5). However, the difference was not statistically significant.

**Figure 2.**
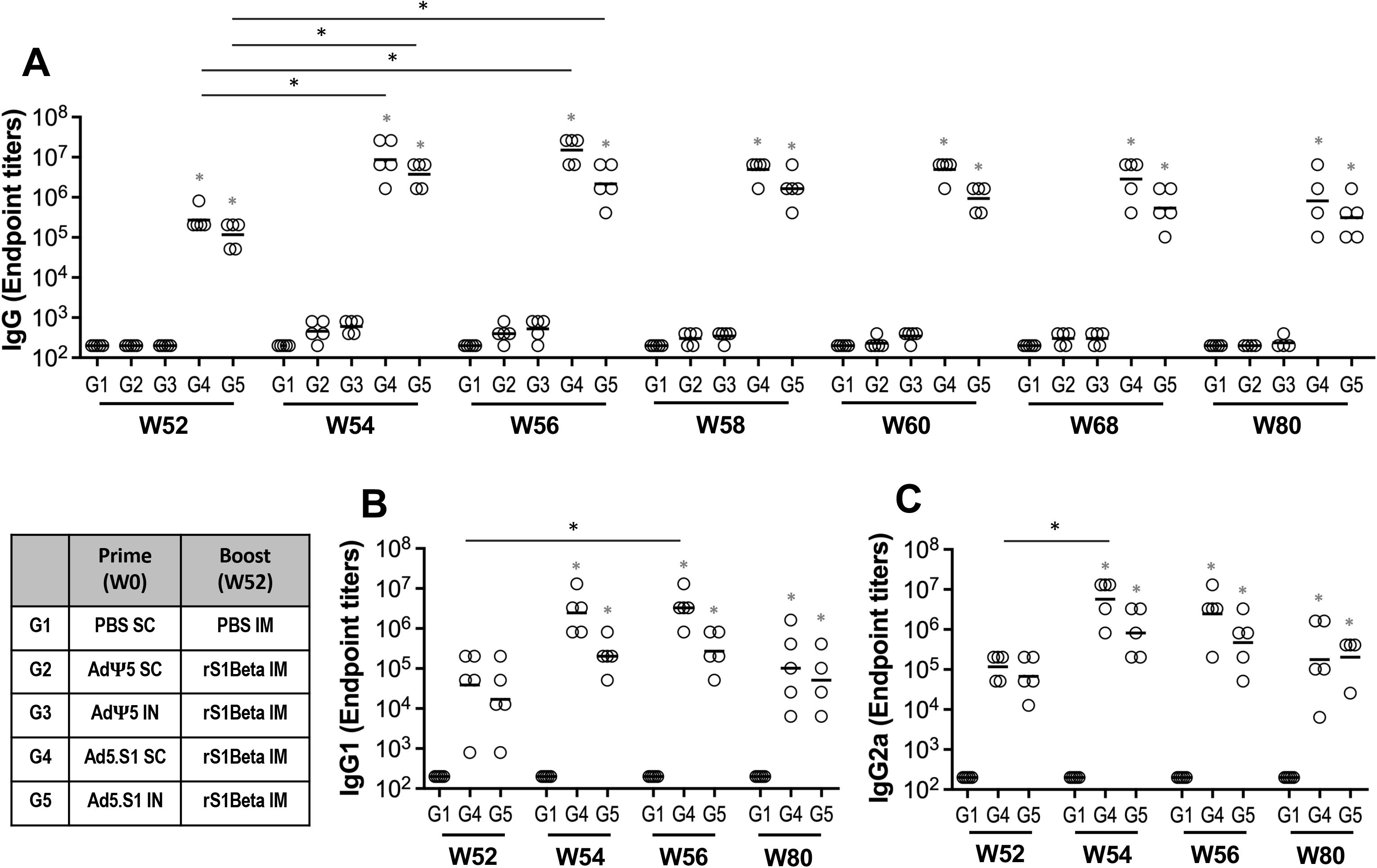
Prime-boost immunization of SARS-CoV-2 adenoviral vaccine-subunit proteins in BALB/c mice. BALB/c mice were primed with 1.5×10^10^ vp of adenoviral vaccine (Ad5.SARS-CoV-2-S1 (Ad5.S1) or AdΨ5) subcutaneously or intranasally, and with PBS as a negative control, and boosted with 15 μg of SARS-CoV-2-S1Beta recombinant proteins intramuscularly at a one-year interval, and immune responses assessed at 52, 54, 56, 58, 60, 68, and 80 weeks post-prime (N=5 per group, except G4 at week 80 N=4). Reciprocal serum endpoint dilutions of SARS-CoV-2-S1-specific antibodies were measured by ELISA to determine the **A.** IgG (at weeks 52, 54, 56, 58, 60, 68, and 80) from all groups, **B.** IgG1 (at weeks 52, 54, 56, and 80) from G1, G4, and G5, and **C.** IgG2a (at weeks 52, 54, 56, and 80) from G1, G4, and G5. Horizontal lines represent geometric mean antibody titers (GMT). Significance was determined by Kruskal-Wallis test, followed by Dunn’s multiple comparisons (\**p* < 0.05). Grey asterisks in Fig.2 represented statistical differences compared with G1, PBS group.

Serum samples collected at week 52, 54, 56, an 80 were serially diluted to determine SARS-CoV-2-S1-specific IgG1 and IgG2a endpoint titers for each immunization group, indicating a Th2- or Th1-like response, respectively, using ELISA **(Fig. 2B and 2C)**. As shown in Fig. 2B and 2C, the induction of S1-specific IgG1 and IgG2a antibodies were significant and similar in G4 and G5 after a booster shot, indicating a balanced Th1/Th2 response. Although there were no significant differences of S1-specific IgG1 and IgG2a responses at week 52 compared to G1, more significantly different IgG1 and IgG2a responses were observed in G4 (*p* < 0.001 at weeks 54 and 56; *p* < 0.05 at week 80) than those in G5 (*p* < 0.05 at weeks 54, 56, and 80), when compared with G1. Interestingly, IgG2a (Th1) responses were recalled faster than IgG1 (Th2) in both G4 and G5 (peak at week 54 vs. week 56, respectively). Results suggest that a booster immunization with rS1Beta subunit vaccine induced significantly increased S1-specific IgG, IgG1, and IgG2a endpoint titers, which were recalled quickly **(Figs. 2A-C,** *p* < 0.05, Kruskal-Wallis test, followed by Dunn’s multiple comparisons). Furthermore, the elicited IgG, IgG1, and IgG2a antibody responses remained significantly high with respect to control groups through week 80 (maximum length of the study to date) than after a prime **(Fig.2 and Supplementary Fig.1)**. Together, these results suggest that a booster was capable of generating robust, balanced, and long-lived S1-specific antibody responses in aged mice primed with Ad5.S1 via either S.C. delivery or I.N. administration one year ago.

### Neutralizing Antibody Levels after a Booster

To evaluate the presence of long-term and booster-generated SARS-CoV-2-specific neutralizing antibodies, we used a microneutralization assay (VNT_90_) by testing the ability of sera from immunized mice to neutralize the infectivity of SARS-CoV-2 Wuhan, Beta (B.1.351), and Delta (B.1.617.2) variants and the results were shown in Fig. 3A. SARS-CoV-2-neutralizating antibodies were detected in Ad5.S1 vaccinated mouse groups (G4 and G5) even after one year of vaccination as compared to PBS group (G1) with no significant differences. The geometric mean titers (GMT) of VNT_90_ in G4 and G5 were 33.7 and 28.6 against Wuhan, 20.5 and 31.8 against Beta (B.1.351), and 8.7 and 10.8 against Delta at week 52, respectively. This result clearly showed the low neutralization against Delta (B.1.617.2) among the variants.

**Figure 3.**
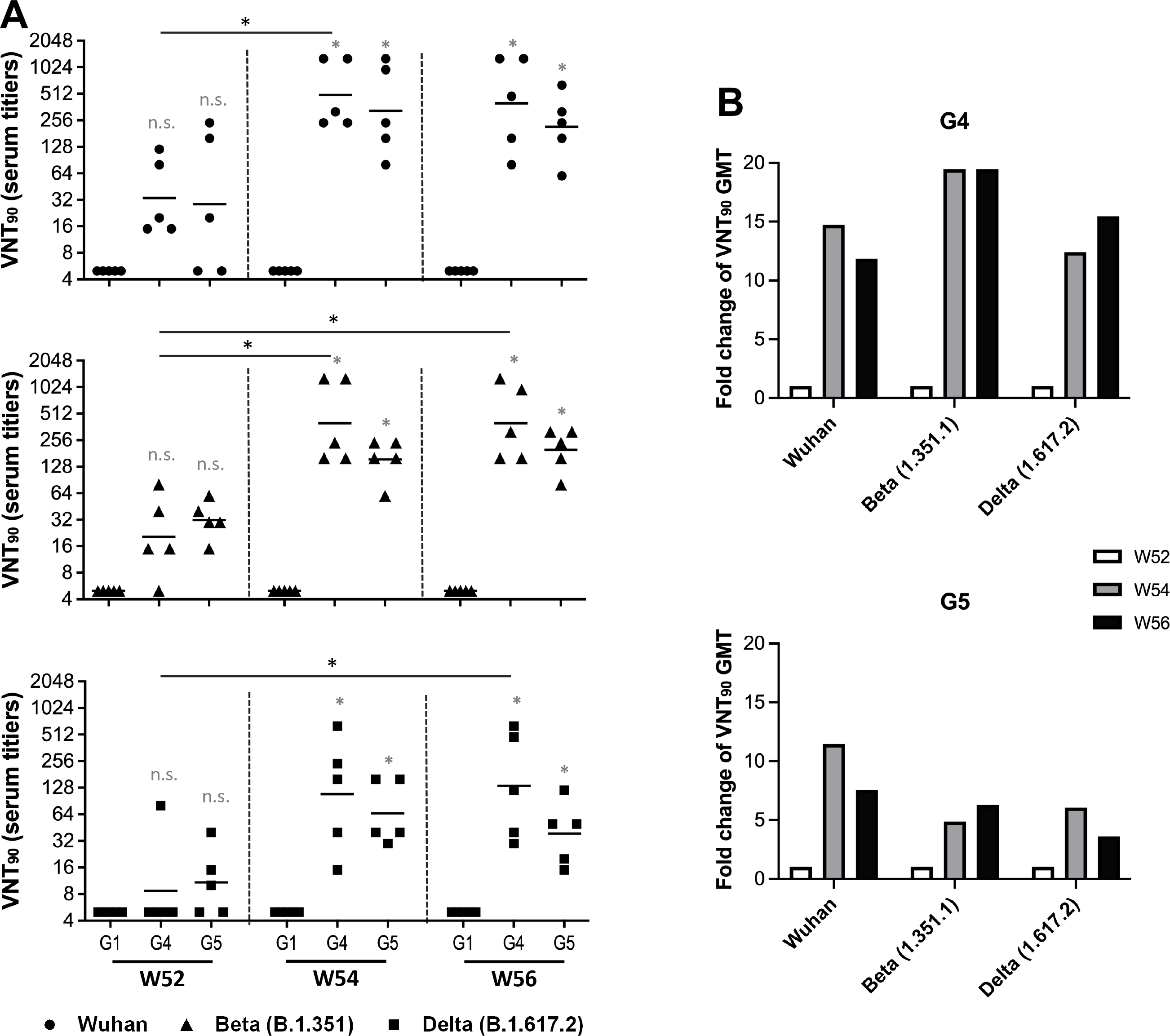
Neutralizing antibody responses in mice after a boost. BALB/c mice (n= 5 mice per group) were prime-immunized subcutaneously or intranasally with 1.5×10^10^ vp of Ad5.SARS-CoV-2-S1 (Ad5.S1) or AdΨ5, respectively, while mice were immunized subcutaneously with PBS as a negative control and boosted with 15 μg of SARS-CoV-2-S1Beta recombinant proteins intramuscularly at week 52. **A.** Neutralizing antibody titers against Wuhan (circle), Beta (B.1.351, triangle), and Delta (B.1.617.2, square) variants of SARS-CoV-2 were measured using a microneutralization assay (VNT_90_) at weeks 52, 54, and 56 after prime immunization. Serum titers that resulted in a 90% reduction in cytopathic effect compared to the virus control were reported. Horizontal lines represent geometric mean neutralizing antibody titers. Groups were compared by Kruskal-Wallis test at each time point, followed by Dunn’s multiple comparisons. Significant differences relative to the PBS control are indicated by *p < 0.05. The minimal titer tested was 10, and undetectable titers (those with NT_90_ serum titers < 10) were assigned a value of 5. Grey asterisks represented statistical differences compared with PBS group. **B.** Fold change of VNT_90_ GMT against Wuhan, Beta (B.1.351), and Delta (B.1.617.2) in G4 and G5 after a booster (weeks 54 and 56, grey and black box, respectively), relative to those of pre-booster (week 52, white box).

After a booster vaccination, the resulting SARS-CoV-2-neutralizing activities on week 54 and week 56 were statistically significant (**Fig. 3A,** p < 0.05, Kruskal-Wallis test, followed by Dunn’s multiple comparisons) compared to control groups, with no significant differences with respect to each other. The fold change of geometric mean titers (GMT) of VNT_90_ against Wuhan ; Beta (B.1.351) ; Delta (B.1.617.2) in G4 compared to those at week 52 were 14.7− ; 19.5− ; 12.4-fold at week 54, and 11.8− ; 19.5− ; 15.5-fold at week 56, respectively (**Fig. 3B)**. Those from G5 were 11.5− ; 4.9− ; 6.1-fold at week 54, and 7.6− ; 6.3− ; 3.6-fold at week 56, respectively. These fold changes of VNT_90_ GMT were statistically significant in G4 against all variants, with no significant differences in G5. Interestingly, the highest fold change was against Beta (B.1.351) in G4, while it was against Wuhan in G5. There were no detected neutralizing antibody responses in the sera from mice immunized with AdΨ5-vaccinated groups (G2 and G3) after a booster (data not shown.)

To assess correlations between levels of S1-binding IgG endpoint titers and levels of neutralizing antibodies, we performed correlation analyses on log-transformed data. We found a positive correlation between S1-binding IgG titers and VNT_90_ in all animals from G1, G4 and G5 at week 52, 54, and 56 (Spearman’s correlation coefficients, *r* = 0.9177 (95%CI: 0.8462-0.9567) for Wuhan, *r* = 0.9498 (95%CI: 0.9047-0.9738) for Beta, *r* = 0.8875 (95%CI: 0.7925-0.9404) for Delta, *p* <0.0001) (**Supplementary Fig.2**). The highest to lowest correlation between S1-binding IgG endpoint titers and neutralizing antibodies was Beta, Wuhan, and Delta, respectively, remarked Beta was a variant of subunit vaccine as a booster.

### ACE2 binding inhibition

Additional tests to evaluate the ability of antibodies in serum were conducted by measuring the inhibition of binding between angiotensin converting enzyme-2 (ACE2) and trimeric spike protein of SARS-CoV-2 variants. We used V-PLEX SARS-CoV-2 (ACE2) Kit Panel 18 including Wuhan, Alpha (B.1.1.7), Beta (B.1.351), Gamma (P.1), Delta (B.1.617.2), Zeta (P.2), Kappa (B.1.617.1), New York (B.1.516.1), India (B.1.617 and B.1.617.3). Antibodies capable of neutralizing the interaction between spike of SARS-CoV-2 variants and ACE2 were examined in all animals from G4 and G5 at week 0, 6, 28, 54, and 80 (**Fig. 4**). The ACE2 inhibitory activities of the antibodies from G4 against all variants were on average 13.2% ± 6.98, 13.3% ± 6.83, 94.9% ± 6.80, and 52.9% ± 36.47 at week 6, 28, 54, and 80, respectively, and those from G5 were on average 14.7% ± 4.82, 14.7% ± 10.87, 74.1% ± 25.38, and 25.2% ± 18.11, respectively with 6.4% ± 2.65 at week 0. Overall, the median percent inhibition was lower for all variants compared to Wuhan wild type. Interestingly, the significant difference for all variants reached statistically in both G4 and G5 groups at week 54, when compared to week 0 (**Fig. 4A and 4B**). The inhibition against Wuhan and Alpha (B.1.1.7) spike by vaccine-induced antibodies at week 80 was significantly different compared to week 0 in only G4 (**Fig. 4A and 4B**). The increase and decrease in percent inhibition towards the different variants followed the same trend for both groups. The highest and lowest percent inhibition of neutralizing antibodies compared to Wuhan was Alpha (B.1.1.7) and Delta (B.1.617.2), respectively.

**Figure 4.**
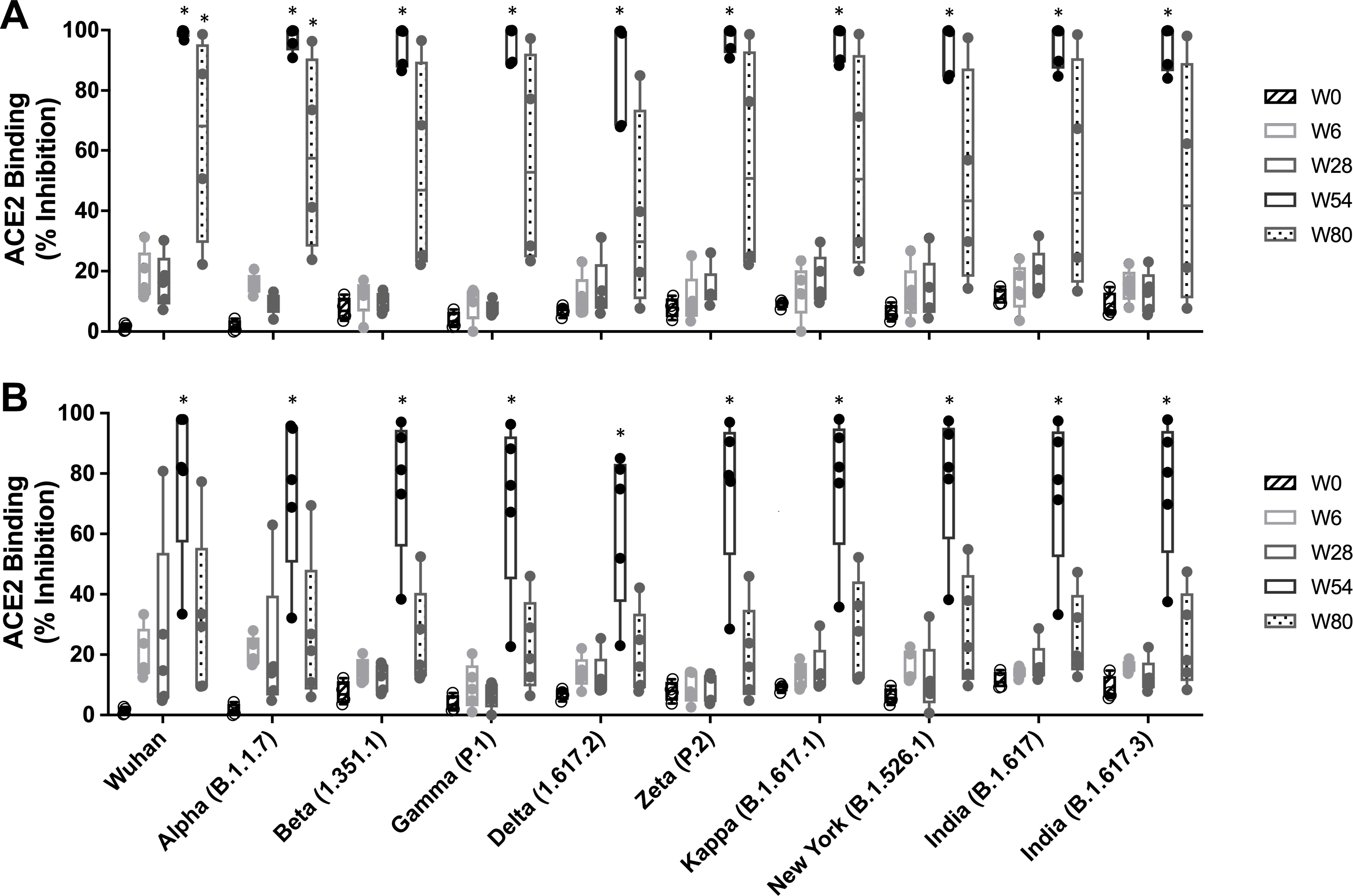
Percent ACE binding inhibition of neutralizing antibodies against SARS-CoV-2 variants. Antibodies in sera capable of neutralizing the interaction between SARS-CoV-2 Wuhan, Alpha (B.1.1.7), Beta (B.1.351), Gamma (P.1), Delta (B.1.617.2), Zeta (P.2), Kappa (B.1.617.1), New York (B.1.516.1), India (B.1.617 and B.1.617.3) variants spike and ACE2 were examined in all animals from G4 **(A)** and G5 **(B)** at week 0 (black with stripes), 6 (light grey), 28 (dark grey), 54 (black), and 80 (dark grey with spots). Serum samples were diluted in 1:100 before adding the V-PLEX plates. Box and whisker plots represent the median and upper and lower quartile (box) with min and max (whiskers). There is no significance difference among all the variants at same time points, neither before, nor after a booster. Grey asterisks represented statistical differences compared with preimmunized sera.

After a booster, ACE2 binding inhibition and VNT_90_ increased significantly against Wuhan, Beta (B.1.351), and Delta (B.1.617.2) compared to the pre-vaccinated sera, with no difference found among the variants. To determine correlations between levels of ACE2 inhibition and levels of neutralizing antibodies, we performed correlation analyses on ACE2 inhibition of 1:100 diluted mice sera and log transformed VNT_90_ data of Wuhan, Alpha (B.1.1.7), and Delta (B.1.617.2). We found a positive correlation between V-PLEX ACE2 inhibition and VNT_90_ in all animals from G1, G4 and G5 at week 54 (Spearman’s correlation coefficients, *r* = 0.9025 (95%CI: 0.8190-0.9486, *p* <0.0001) (**Fig. 5**). Spearman’s correlation coefficients were lower when analysis was performed with 1:400 diluted mice sera (*r* = 0.7802 (95%CI: 0.6132-0.8804, *p* <0.0001) (**Supplementary Fig. 3**). Changes of ACE2 binding inhibition at week 6, 28, 54, and 80 against Wuhan spike protein were dependent on dilution factor, showing the similar pattern with other variants (**Supplementary Fig. 1C**). Taken together, single dose of non-adjuvanted recombinant S1 protein subunit vaccine as a booster induced robust and broadly cross-reactive neutralizing antibodies against SARS-CoV-2 variants in aged mice and neutralizing antibody titer was correlated with inhibition of spike-ACE2 binding positively.

**Figure 5.**
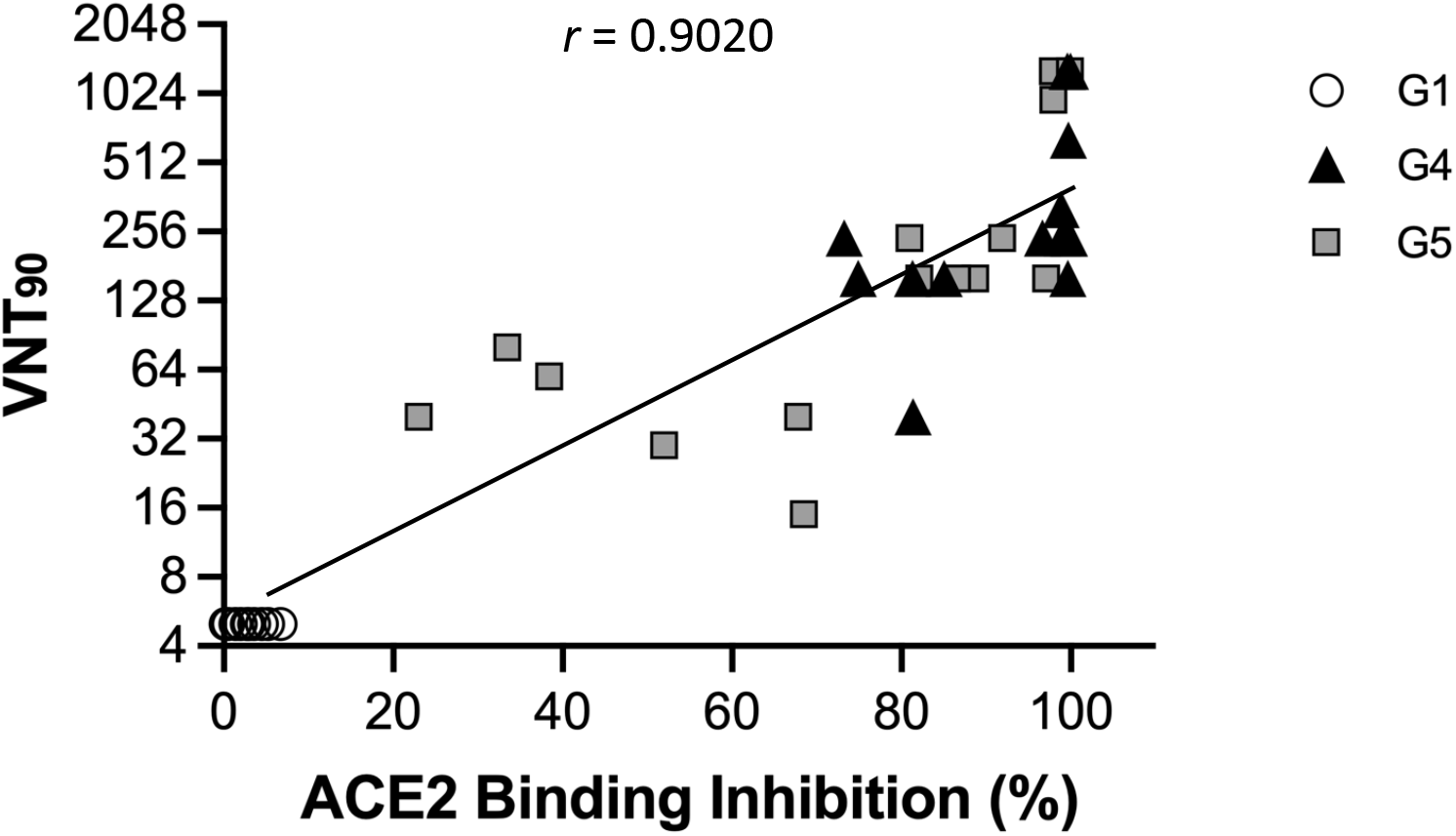
Correlation between the VNT_90_ and ACE2 binding inhibition. Correlation between VNT_90_ (Log_2_) against Wuhan, Beta (B.1.351), and Delta (B.1.617.2) and ACE2 binding inhibition (%) of 1:100 diluted sera from all animals from G1 (white circle), G4 (black triangle), and G5 (grey square) at week 54. The lines represent the regression line of all samples. Each symbol represents an individual mouse. Correlation analysis and calculation of Spearman’s correlation coefficients was performed using GraphPad Prism v9.

## Discussion

We previously reported that a single immunization of BALB/c mice via either I.N. or S.C. delivery of our adenovirus-based COVID-19 vaccine (Ad5.S1) elicited robust S1-specific humoral and cellular immune responses in mice. In this study, we demonstrated the long-term persistence of immunogenicity after prime vaccination until one year. We also demonstrated that a booster of non-adjuvanted recombinant S1 proteins of Beta variant induced a robust balanced long-lasting IgG antibodies and neutralizing antibodies, which were broadly cross-reacting with SARS-CoV-2 variants and corelated with ACE2-spike interaction inhibition.

There were very low antibody responses in the sera from mice immunized with AdΨ5-vaccinated groups (G2 and G3) after a subunit booster injection at week 52, which might be explained by the age of the mice at the time of single immunization **(Fig. 2A).** Indeed, vaccinated aged mice elicited lower level of immune responses compared to vaccinated young mice, which were found to be due to a low frequency of IgG- and IFN-*γ*-secreting cells in vaccinated aged mice [33]. These results were parallel to previous findings that older individuals have lower immune responses to approved COVID-19 vaccine than younger individuals [10, 12, 33, 34]. Specially, lower serum IgG levels of SARS-CoV-2 in elderly peoples were from a lower proportion of peripheral spike-specific memory B cells [12].

Although there was no significant differences between mice groups primed with Ad5.S1 SC or IN, animal group primed SC showed long lasting and higher GMT after a booster than animal group primed IN **(Fig. 2A)**. However, it may not guarantee that SC injection will be better than IN delivery for protection against existing and newly emerging SARS-CoV-2 variants. Various studies reported that vaccine delivered intranasally elicited superior mucosal immunity compared to the intramuscular injection, and protected efficiently after the challenge and reduced viral transmission [35–38]. Moreover, a recent study of adjuvanted S1 subunit vaccines primed-boosted intramuscularly or primed intramuscularly-boosted intranasally in rhesus macaques reported that the mucosal vaccine demonstrated outstanding protection in both upper and lower respiratory tracts by clearing of the input virus more efficiently through higher dimeric IgA and IFN-α in BAL fluid, although intranasal boosting elicited weaker T cell and lower neutralizing antibody titer [35].

In this study, high titer of serum S1-binding IgG was investigated up to 28 weeks after a booster in aged mice primed Ad5.S1 one year ago **(Supplement Fig1B)**. Although limits of IgG duration of mice may not be reflective of that measured in non-human primates or humans, this result implied that humoral immunity might be long-lasting after a booster, because IgG titers at 28 weeks post-booter in G4 and G5 was approximately 6-fold and 1.7-fold higher than those at 28 weeks post-prime, respectively. Indeed, boosting enhanced humoral and cell-mediated immune responses dramatically in aged mice [34]. Likewise, one of the approved COVID-19 vaccine, Ad26.COV2.S, which is single-shot regimen vaccine protecting against severe COVID-19, induced durable immune responses detected up to 8 months after vaccination in human, although it remains unclear how long immune protection will last after previous infection or vaccination due to the limited length of follow-up studies [39]. The protection of two doses of mRNA BNT162b2 vaccine waned considerably after 6 months in human. However, infection-acquired immunity boosted with vaccination remained high for more than 1 year after infection [40].

Subunit vaccine booster elicited both of high S1-specific IgG1 and IgG2a subclass antibodies in aged mice primed with Ad5.S1, indicating a balanced Th1/Th2 response (**Fig. 2B and 2C**), whereas subunit vaccine alone induced high IgG1 with lower IgG2a leading a possibility of vaccine-associated enhanced respiratory disease (VARED) [41]. Indeed, VARED-like pulmonary immunopathology related with Th2-based immune responses was observed in animals with whole-inactivated SARS-CoV vaccines [42, 43]. In this study, a high level of neutralizing antibodies and the balanced Th1/Th2 immune response were induced, suggesting that a booster of subunit vaccine after an adenoviral prime vaccine might avoid Th2-based immune response and the occurrence of VAERD.

Neutralization assay was frequently used as a correlate of protection following vaccination [24–27, 44]. Here we used a microneutralization test to evaluate the function of the generated antibodies in the sera of immunized mice. The titer of neutralizing antibodies dramatically increased after a booster and neutralized other variants of Beta and Gamma (**Fig 3**). Our future studies will include the evaluation of neutralization effect against omicron variant. Notably, a recent study demonstrated that the boosted immune response by mRNA BNT162b2 neutralized omicron variant [34]. If needed, it may be possible to further improve neutralizing antibody responses with a booster of omicron BA.5 rS1 subunit vaccine to overcome the emerging SARS-CoV-2 infection. Neutralizing antibodies against SARS-CoV-2 are effective at blocking spike-ACE2 binding to prevent infection [16, 17]. As a conventional pseudo-neutralizing test, measurement of competitive immunoassay for quantifying inhibition of the spike-ACE2 interaction can be used as a surrogate for traditional virus-based plaque reduction neutralizing assay and reported in a high level of concordance and correlation (>96%) [18, 19]. In this study, we assessed animal immune response for blocking spike-ACE2 binding using V-PLEX neutralization panel kit and showed that a booster of aged mice primed with Ad5.S1 could induce significant blocking in binding of ACE2 to spike of all tested variants (**Fig 4**), which was correlated with VNT_90_ (**Fig 5**). In addition, our future work will include more investigation in blocking of binding of ACE2 to spike of omicron variants.

Here we showed non-adjuvated subunit vaccine booster effect. However, there will be a beneficial effect of an adjuvanted subunit booster strategy for protection, especially against distant variants such as Omicron BA.5. Actually, AS03-adjuvanted CoV2 preS dTM (B.1.351) induced higher neutralizing antibody titers against Beta variant in non-human primates compared to animal group boosted with non-adjuvanted vaccine in the mRNA-primed cohort [21]. AS01-like adjuvanted SARS-CoV-2 subunit vaccine enhanced Th1 type-IgG2a isotype, neutralizing antibodies, and IFN-γ secreting T cell immune responses in both young and aged mice [45]. Recombinant S protein in combination with adjuvant CoVaccine HT™ induced a balanced IgG subtype antibody response [41].

Two limitations of this study were T-cell immunity and SARS-CoV-2 challenge, which were not performed to test the cellular immunity and to assess the protection efficiency of booster vaccination. However, various studies have reported previously that T-cell immunity was activated after a booster [23, 35, 46, 47]. Homologous and heterologous boosters in health care workers who had received a priming dose of Ad26.COV2.S Covid-19 vaccine resulted in higher levels of T-cell responses than non-booster group, although T-cell response was significantly larger with mRNA-based vaccines (91%) than with the homologous booster (72%) [23]. Additionally, a booster dose of mRNA BNT162b2 elicits robust T cell responses that cross-recognized SARS-CoV-2 Omicron variant in aged mice [34]. Not only mRNA vaccine but also adenoviral vector or adjuvanted protein subunit vaccines enhanced cellular immune response in aged mice after a boost [13, 48]. Furthermore, S-specific T-cell responses were positively correlated with the presence of S-specific binding antibodies [23], implying induction of robust T cell immune response after rS1beta booster in this study.

As our study does not define protection ability against SARS-CoV-2 variants by challenge, it needs to be investigated in the future. Notably, a recent study was performed a protection experiment against SARS-CoV-2 Omicron variant in aged BALB/c mice boosted with mRNA vaccine [34]. This natural mouse model of SARS-CoV-2 infection by assessing viral replication and histopathological changes in the lung, does not require genetic modification of mice or viruses. However, this wild mouse animal model only supports infection of SARS-CoV-2 variants that carry the N501Y mutation, including Alpha, Beta, Gamma, and Omicron [49]. Therefore, it is still important that K18-hACE2 and other hACE2-transgenic mice are also used to investigate pathogenicity of different SARS-CoV-2 variants [50].

Overall, our study evaluated the effect of a booster in aged mice after priming of adenoviral vaccines as a pre-clinical model of elderly peoples immunized with the current approved Covid-19 vaccines. Our findings may have implications for further study of using recombinant protein S1BA.5 subunit vaccine as a booster to enhance cross-neutralizing antibodies against new emerging variants of concern.

## Materials and methods

### Construction of recombinant protein expressing vectors

The coding sequence for SARS-CoV-2-S1 amino acids 1 to 661 [51] mutated at del144; K417N; E484K; N501Y; A570D; D614G having C-terminal tag known as ‘C-tag’, composed of the four amino acids (aa), glutamic acid–proline–glutamic acid–alanine (E-P-E-A) flanked with Sal I & Not I was codon-optimized using the UpGene algorithm for optimal expression in mammalian cells [52] and synthesized (GenScript). The construct also contained a Kozak sequence (GCCACC) at the 5′ end. The plasmid, pAd/SARS-CoV-2-S1Beta was then created by subcloning the codon-optimized SARS-CoV-2-S1Beta inserts into the shuttle vector, pAdlox (GenBank U62024), at Sal I/Not I sites. The plasmid constructs were confirmed by DNA sequencing.

### Transient Production in Expi293 Cells

pAd/SARS-CoV-2-S1Beta was amplified and purified using ZymoPURE II plasmid maxiprep kit (Zymo Research). For Expi293 cell transfection, we used ExpiFectamie™ 293 Transfection Kit (ThermoFisher) and followed the manufacturer’s instructions. Cells were seeded 3.0 × 10^6^ cells/ml one day before transfection and grown to 4.5~5.5 × 10^6^ cells/ml. 1μg of DNA and ExpiFectamine mixtures per 1ml culture were combined and incubated for 15 min before adding into 3.0 × 10^6^ cells/ml culture. At 18-22h post-transfection, enhancer mixture was added, and culture was shifted to 32°C. The supernatants were harvested at 5 days post transfection and clarified by centrifugation to remove cells, filtration through 0.8μm, 0.45μm, and 0.22μm filters and either subjected to further purification or stored at 4°C before purification.

### SDS-PAGE and western blot

To evaluate the expression of the constructed plasmids, Expi293 cells were transfected with pAd/SARS-CoV-2-S1Beta. At 5 days after transfection, the supernatants were subjected to sodium dodecyl sulfate polyacrylamide gel electrophoresis (SDS-PAGE) and Western blot as previously described [1, 51]. Briefly, after blocking, rabbit anti-SARS-CoV-2 spike polyclonal antibody (1:3000, Sino Biological) was added and incubated overnight at 4 °C as primary antibody, and horseradish peroxidase (HRP)-conjugated goat anti-rabbit IgG (1:10000, Jackson immunoresearch) was added and incubated at RT for 2 hours as secondary antibody. After washing three times with PBST, the signals were visualized on an iBright FL 1500 Imager (ThermoFisher).

### Purification of recombinant proteins

The recombinant proteins named rS1Beta was purified using a CaptureSelect™ C-tagXL Affinity Matrix prepacked column (ThermoFisher) and followed the manufacturer’s guidelines. Briefly, The C-tagXL column was conditioned with 10 column volumes (CV) of equilibrate/wash buffer (20 mM Tris, pH 7.4) before sample application. Supernatant was adjusted to 20 mM Tris with 200 mM Tris (pH 7.4) before being loaded onto a 5-mL prepacked column per the manufacturer’s instructions with 5 ml/min rate. The column was then washed by alternating with 10 CV of equilibrate/wash buffer, 10 CV of strong wash buffer (20 mM Tris, 1 M NaCl, 0.05% Tween-20, pH 7.4), and 5 CV of equilibrate/wash buffer. The recombinant proteins were eluted from the column by using elution buffer (20 mM Tris, 2 M MgCl_2_, pH 7.4). The eluted solution was concentrated and desalted with preservative buffer (PBS) in an Amicon Ultra centrifugal filter devices with a 50,000 molecular weight cutoff (Millipore). The concentration of the purified recombinant proteins was determined by the BCA protein assay kit (Thermo Scientific) using bovine serum albumin (BSA) as a protein standard, separated by reducing SDS-PAGE, and visualized by silver staining.

### Animals and immunization

At week 52, female BALB/c mice (n = 5 animals per group) primed with adenovirus-based COVID-19 vaccine (Ad5.S1) [1] were boosted by intramuscularly with 15 μg of rS1Beta in the thigh or PBS as a negative control. Mice were bled from retro-orbital vein at weeks 52, 54, 56, 58, 60, 68, and 80 after prime immunization, and the obtained serum samples were diluted and used to evaluate S1-specific antibodies by enzyme-linked immunosorbent assay (ELISA). Serum samples obtained on weeks 52, 54, and 56 were also used for microneutralization (NT) assay. Mice were maintained under specific pathogen-free conditions at the University of Pittsburgh, and all experiments were conducted in accordance with animal use guidelines and protocols approved by the University of Pittsburgh’s Institutional Animal Care and Use (IACUC) Committee.

### ELISA

To evaluate the expression of SARS-CoV-2S1Beta recombinant protein, ELISA plates were coated with chimeric MAb 40150-D003 (1:750, Sino Biological) overnight at 4°C in carbonate coating buffer (pH 9.5) and then blocked with PBS containing 0.05% Tween 20 (PBS-T) and 2% bovine serum albumin (BSA) for one hour. The supernatants of Expi293™ cells transfected with pAd/SARS-CoV-2-S1SA was diluted 1:40 in PBS-T with 1% BSA and along with standard control protein 40591-V08H (rS1H, Sino Biological) or purified rSARS-CoV-2S1Beta were incubated overnight at 4°C. After the plates were washed, chimeric MAb 40150-D001 HRP conjugated secondary antibody (1:10000, Sino Biological) was added to each well and incubated for one hour. The plates were than washed three times and developed with 3,3’5,5’-tetramethylbenzidine, and the reaction was stopped with 1M H_2_SO_4_ and absorbance at 450 nm was determined using an ELISA reader (Molecular Devices SPECTRAmax).

To investigate the immunogenicity of SARS-CoV-2S1Beta recombinant protein, ELISA was performed as previously described [1, 51, 53]. Sera from all mice were collected prior to boost (week 52) and every two weeks (week 54, 56, 58, 60) after immunization and tested for SARS-CoV-2-S1-specific IgG, IgG1, and IgG2a antibodies using conventional ELISA. Furthermore, sera from all mice collected at weeks 68 and 80 after prime immunization were tested for SARS-CoV-2-S1-specific IgG antibodies using ELISA for long-term humoral responses. Sera collected at week 28 (W80) after boost vaccination were also tested for SARS-CoV-2-S1-specific IgG1 and IgG2a antibodies using ELISA. Briefly, ELISA plates were coated with 200 ng of recombinant SARS-CoV-2-S1 protein (Sino Biological) per well overnight at 4°C in carbonate coating buffer (pH 9.5) and then blocked with PBS containing 0.05% Tween 20 (PBS-T) and 2% bovine serum albumin (BSA) for one hour. Mouse sera were diluted in PBS-T with 1% BSA and incubated overnight. After the plates were washed, anti-mouse IgG-horseradish peroxidase (HRP) (1:10000, Jackson Immunoresearch), or biotin-conjugated IgG1 and IgG2a (1:1000, eBioscience) was added to each well and incubated for one hour. The plates were washed three times and developed with 3,3’5,5’-tetramethylbenzidine, and the reaction was stopped and absorbance at 450 nm was determined using an ELISA reader (Molecular Devices SPECTRAmax).

### SARS-CoV-2 microneutralization assay

Neutralizing antibody (NT-Ab) titers against SARS-CoV-2 were defined according to the following protocol [54, 55]. Briefly, 50 µl of sample from each mouse, starting from 1:10 in a twofold dilution, were added in two wells of a flat bottom tissue culture microtiter plate (COSTAR, Corning Incorporated, NY 14831, USA), mixed with an equal volume of 100 TCID_50_ of a SARS-CoV-2 Wuhan, Beta, or Delta strain isolated from symptomatic patients, previously titrated, and incubated at 33°C in 5% CO_2_. All dilutions were made in EMEM (Eagle’s Minimum Essential Medium) with addition of 1% penicillin, streptomycin and glutamine and 5 γ/mL of trypsin. After 1 hour incubation at 33°C 5% CO_2_, 3×10^4^ VERO E6 cells [VERO C1008 (Vero 76, clone E6, Vero E6); ATCC^®^ CRL-1586™] were added to each well. After 72 hours of incubation at 33°C 5% CO_2_ wells were stained with Gram’s crystal violet solution (Merck KGaA, 64271 Damstadt, Germany) plus 5% formaldehyde 40% m/v (Carlo ErbaSpA, Arese (MI), Italy) for 30 min. Microtiter plates were then washed in running water. Wells were scored to evaluate the degree of cytopathic effect (CPE) compared to the virus control. Blue staining of wells indicated the presence of neutralizing antibodies. Neutralizing titer was the maximum dilution with the reduction of 90% of CPE. A positive titer was equal or greater than 1:10. The geometric mean titers (GMT) of VNT_90_ end point titer were calculated with 5 as a negative shown <10. Sera from mice before vaccine administration were always included in microneutralizaiton (VNT) assay as a negative control.

### ACE2 blocking assay

Antibodies blocking the binding of SARS-CoV-2 spike variants (Alpha (B.1.1.7), Beta (B.1.351), Gamma (P.1), Delta (B.1.617.2), Zeta (P.2), Kappa (B.1.617.1), New York (B.1.516.1), India (B.1.617 and B.1.617.3)) to ACE2 were detected with a V-PLEX SARS-CoV-2 Panel 18 (ACE2) Kit (Meso Scale Discovery (MSD)) according to the manufacturer’s instructions. The assay plate was blocked for 30 min and washed. Serum samples were diluted (1:25, 1:100 or 1:400) and 25 μl were transferred to each well. The plate was then incubated at room temperature for 60 min with shaking at 700 rpm, followed by the addition of SULFO-TAG conjugated ACE2, and continued incubation with shaking for 60 min. The plate was washed, 150 μl MSD GOLD Read Buffer B was added to each well, and the plate was read using the QuickPlex SQ 120 Imager. Electrochemiluminescent values (ECL) were generated for each sample. Results were calculated as % inhibition compared to the negative control for the ACE2 inhibition assay, and % inhibition is calculated as follows: % neutralization = 100 × (1 − (sample signal/negative control signal)).

### Statistical analysis

Statistical analyses were performed using GraphPad Prism v9 (San Diego, CA). Antibody endpoint titers and neutralization data were analyzed by Kruskal-Wallis test, followed by Dunn’s multiple comparisons. Significant differences are indicated by * p < 0.05. Comparisons with non-significant differences are not indicated. Correlations between the V-PLEX ACE2 blocking and VNT_90_ or IgG end point titers and VNT_90_ were determined using correlation analysis and calculation of Spearman coefficients and 95% confidence interval (95% CI).

---

**Supplementary Figure 1.** Comparison of serological responses to S1 from mice of G1, G4, and G5 post-prime and post-boost. Mice were primed with adenoviral vaccine subcutaneously or intranasally and boosted with SARS-CoV-2-S1Beta recombinant proteins at a one-year interval, and reciprocal serum endpoint dilutions of S1- specific IgG were measured by ELISA. **A.** Fold change of reciprocal serum endpoint dilutions of S1-specific IgG in G4 (white box) and G5 (grey box) after a booster compared to those at week 52 **B.** Serum endpoint titers of S1- specific IgG were assessed at highest time point (weeks 6 and 54) and at week 28 post-prime or post-boost (N=5 per group, except at week 80 G4 N=4). Horizontal lines represent geometric mean antibody titers (GMT). Significance was determined by Kruskal-Wallis test, followed by Dunn’s multiple comparisons (\**p* < 0.05). **C.** ACE2 binding inhibition (%) at weeks 6, 28, 54, and 80 against Wuhan in dilution 1:25, 1:100, and 1:400. Data showed means ± standard error of the means (SEM) of G1 (white circle), G4 (black triangle), and G5 (grey square).

**Supplementary Figure 2.** Correlation between the VNT_90_ and SARS-CoV-2-S1-specific IgG titer. Correlation between VNT_90_ (Log_2_) against Wuhan, Beta (B.1.351), and Delta (B.1.617.2), and S1-binding IgG endpoint titers (Log_10_) in all animals from G1 (white circle), G4 (black triangle), and G5 (grey square) at week 52, 54, and 56. The lines represent the regression line of all samples. Each symbol represents an individual mouse. Correlation analysis and calculation of Spearman’s correlation coefficients was performed using GraphPad Prism v9.

**Supplementary Figure 3.** Correlation between VNT_90_ (Log_2_) against Wuhan, Beta (B.1.351), and Delta (B.1.617.2) and ACE2 binding inhibition (%) of 1:400 diluted sera from all animals of G1 (open circle), G4 (black triangle), and G5 (grey square) at week 54. The line represents the regression line of all samples. Correlation analysis and calculation of Spearman’s correlation coefficients was performed using GraphPad Prism 9.

## Notes

### Competing Interest Statement

The authors have declared no competing interest.

