## Supplementary figures and images for "SARS-CoV-2 S1 Subunit Booster Vaccination Elicits Robust Humoral Immune Responses in Aged Mice"

### Supplementary Figures

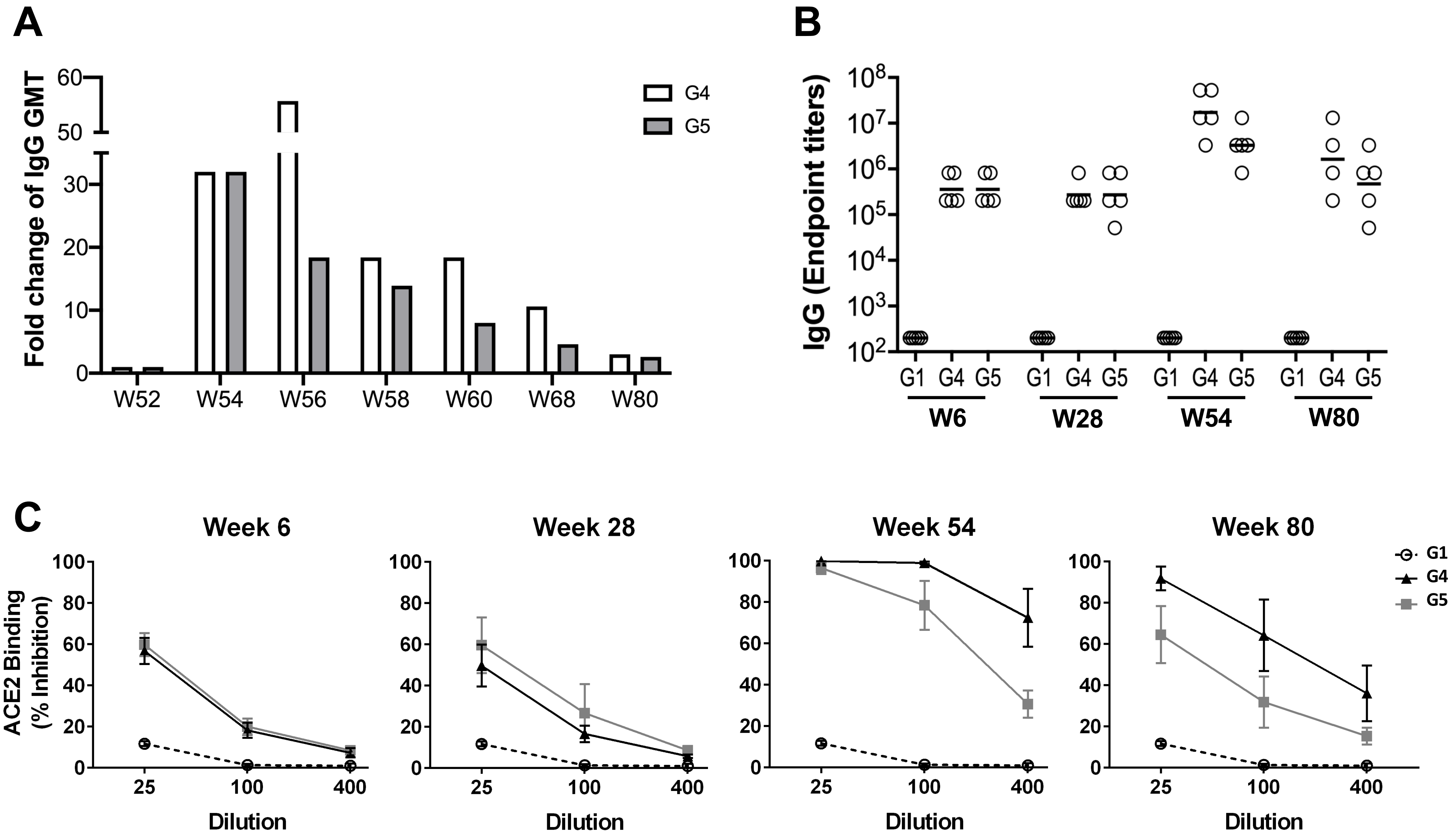


Supplementary Figure 1

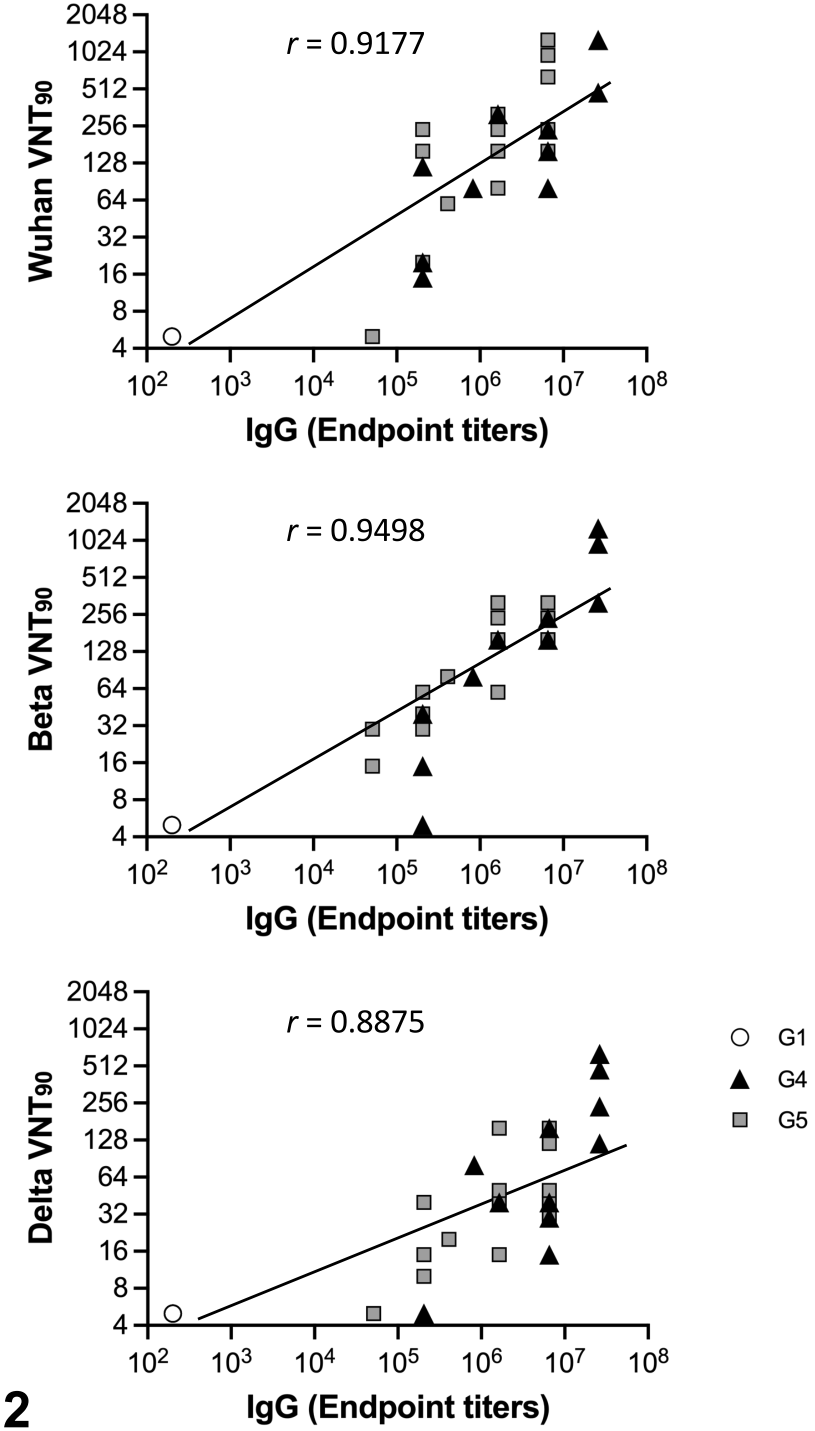


**Supplementary Figure 2**

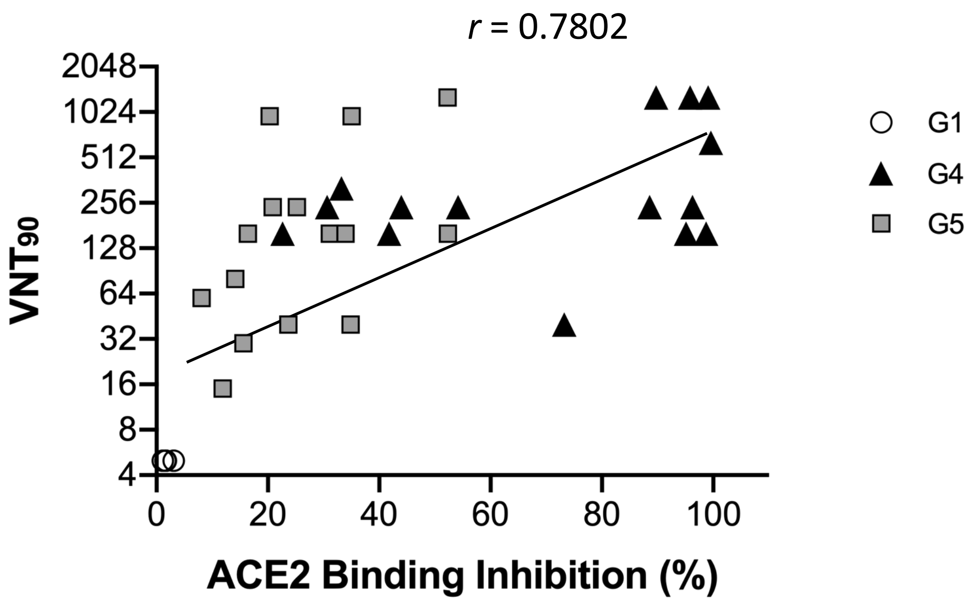


**Supplementary Figure 3**
